# Detecting epidemic-driven selection: a simulation-based tool to optimize sampling design and analysis strategies

**DOI:** 10.1101/2024.06.27.601009

**Authors:** Cindy G. Santander, Ida Moltke

## Abstract

Throughout history, populations from numerous species have been decimated by epidemic outbreaks, like the 19^th^-century rinderpest outbreak in Cape buffalo (*≈* 90% mortality) and Black Death in humans (*≈* 50% mortality). Recent studies have raised the enticing idea that such epidemic outbreaks have led to strong natural selection acting on disease-protective variants in the host populations. However, so far there are few, if any, clear examples of such selection having taken place. This could be because so far studies have not had sufficient power to detect the type of selection an epidemic outbreak must induce: strong but extremely short-term selection on standing variation. We present here a simulation-framework that allows users to explore under what circumstances it is possible to detect epidemic-driven selection using standard selection scan methods like *F_ST_*and iHS. Using two examples, we illustrate how the framework can be used. Furthermore, via these examples, we show that comparing survivors to the dead has the potential to render higher power than more commonly used sampling schemes. And importantly, we show that even for outbreaks with high mortality, like the Black Death, strong selection may have led to only modest shifts in allele frequency, suggesting large sample sizes are required to obtain appropriate power to detect the selection. We hope this framework can help in designing well-powered future studies and thus lead to a clarification of the role epidemic-driven selection has played in the evolution of different species.

**Significance Statement:** Our study introduces a simulation-based framework, `SimOutbreakSelection` (`SOS`), which enables researchers to design studies that have power to detect epidemic-driven selection while taking sampling time points and demographic history into account. We use rinderpest in African Buffalo and the Black Death in Medieval Sweden as examples to showcase the framework. Via these examples we also show that large sample sizes are needed even for severe epidemics like the Black Death and that the often used sampling strategy where samples from before the epidemic and samples from after are compared is not always optimal.

## 1 Introduction

Outbreaks of severe disease epidemics have had devastating effects on many human and animal populations throughout history. Since the worst outbreaks are known to have killed large proportions of the affected populations, any potential genetic variant that increased the ability of its carriers to survive the outbreak must have been under extremely strong positive natural selection. Nevertheless, evidence of selection driven by severe epidemic outbreaks has been limited, despite efforts to investigate the effect on the host population of several outbreaks with high mortality [1].

Early studies delved into selection driven by epidemics on a genic-level [2–5] and recently researchers have begun conducting selection scans to address some of these questions [1, 6–9]. For instance, the Black Death, a canonical example of an epidemic disease, has been extensively studied in both ancient and modern individuals [8–12], yet no clear and undisputed evidence of epidemic-driven selection has been found in these selection scans [7, 9, 13]. It is possible that the lack of wide-spread evidence is due to this type of selection rarely, or never, occurring because it requires a protective variant to already be present in the population when the outbreak takes place. However, it may also simply be due to limitations in the studies that have been performed, such as limited sample size. While the ability to study ancient DNA has expanded the scope of research on selection across time [14–18], the number of samples that have been analysed so far may not have been sufficient to answer questions related to epidemic-driven selection [13]. Moreover, commonly used methods for detecting selection in population genetics are optimized for continuous selection over many generations acting on new variants [19]. In contrast, selection driven by epidemic outbreaks is short-term (the time each outbreak takes) and must necessarily act on standing variation, because the short term nature of outbreaks leaves prohibitively short time for a new advantageous mutation to occur and spread in a population. Lastly, most current methods rely on detecting allele frequency changes that occur only at the selected locus [20]—a notable limitation given the nature of epidemics to strongly increase genetic drift as a consequence of mass death in a population [21].

In light of the aforementioned considerations and limitations, we here introduce a simulation-based framework called `SimOutbreakSelection` (`SOS`), which allows users to explore under what settings (e.g. sample size, sampling scheme, selection scan method)—if any—it is possible to detect epidemic-driven selection on a single advantageous variant in their population and epidemic of interest. For instance, SOS can be used to see if the number of samples one has access to are enough to have reasonable power to detect selection and, if so, which methods would provide the most power. Similarly, one can explore how many samples would be needed for a given sampling scheme and selection scan method of choice by performing simulations with selection using a range of different sample sizes, and for each of these, estimating power as the proportion of times the variant is detected to be under selection. Through this framework, we aim to facilitate the design of well-powered studies that can reveal the extent to which epidemic-driven selection has actually taken place.

In this paper we will first introduce the framework. Then we will illustrate how it can be used via a few examples and finally we will discuss the benefits and limitations of using it. Importantly, besides illustrating how the framework can be used, the examples also reveal two key points about the power to detect epidemic-driven selection: 1) that large sample sizes will often be needed to obtain adequate power because the increase in allele frequency due to selection can be modest even in an outbreak with a mortality of 50% and 2) that comparing survivors to those that have died from the epidemic is worth considering, since it can enhance detection power compared to several other sampling schemes under certain circumstances.

## 2 Results

### 2.1 Overview of the Framework

Our framework, `SOS`, is based on the forward-simulator `SLiM` [22] and it provides a straightforward means of assessing the power of various methods to detect selection driven by an epidemic. With `SOS`, users can flexibly simulate a customized epidemic scenario featuring a specific genetic variant under selection, sample a chosen number of individuals at one or more designated time points, calculate and plot a range of commonly employed selection scan statistics, and summarize the results from multiple simulations to estimate the power of each method expressed as the proportion of simulations in which selection was detected. This means that users can use the framework to help them design well-powered selection studies of historical epidemics, such as for instance rinderpest in African buffalo and Black Death in Medieval Sweden.

To work, SOS needs three types of input (Fig. 1A). The first type of input is data that mimics the relevant population right before the epidemic occurred. This can be obtained by simulating a realistic demographic history informed by the current literature in addition to the user’s knowledge about their organism. The second type of input is information about the course of the epidemic in question. This includes the number of outbreaks and thus population bottlenecks, bottleneck size(s) and bottleneck length(s).

**Figure 1:**
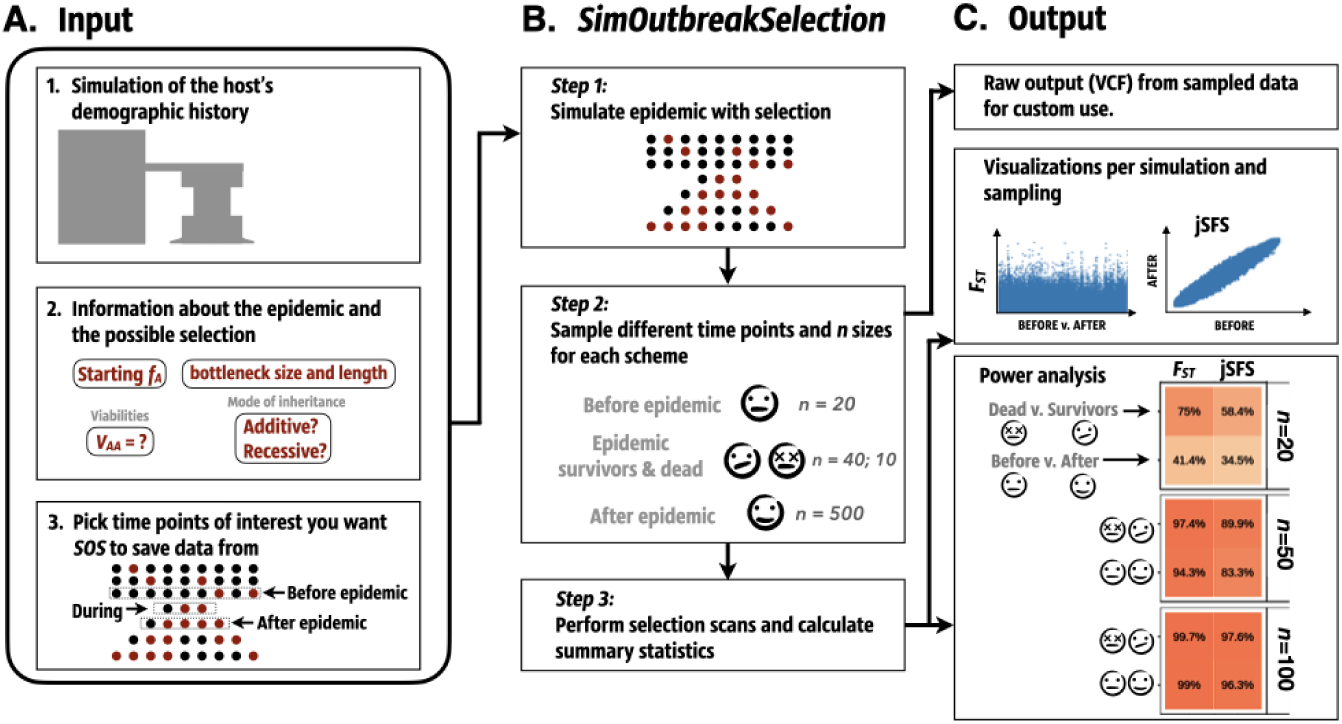
`SOS` Framework. **A. Input:** There are three required input types to use SOS. They are the (1) demographic simulation, (2) the details of the epidemic and (3) the target generations and populations to be saved when simulating the epidemic with selection in SOS. **B.** `SOS`: There are three steps to the `SOS` tool which consists of (1) simulating the epidemic, (2) sampling the different time points and different sample sizes for each sampling scheme, and (3) running different selection scan methods to calculate summary statistics. **C. Output:** `SOS` plots different summary statistics per simulation and sampling. These summary statistics can also be used to calculate power of the different selection scan methods. For convenience, and to make the tool as flexible as possible, `SOS` offers the raw simulation data (i.e. genotypes in a VCF file) of sampled populations and time points as output in case the user wants to explore this in other ways.

Additionally, it includes the user’s assumptions about the potential advantageous variant. For this it is worth noting that to make it possible to sample deceased epidemic victims, the framework is built on the use of a non Wright-Fisher model for the epidemic forward simulations. This also has the benefit that selection acting on the advantageous variant is modelled using viabilities, i.e. a survival probability (or 1-mortality rate) for each genotype (*V_AA_*, *V_Aa_*, *V_aa_*), as opposed to a single relative fitness coefficient, which makes the selection model easier to relate to reported mortality rates during epidemics. For the users, this in practice means that they have to specify the following assumptions about the potential advantageous variant: its allele frequency right before the epidemic (starting frequency), whether its mode of inheritance is additive, recessive or dominant, and the viability of its homozygous carriers (*V_AA_*). Given this—and the size and length of the bottleneck(s)—a fitness model is fully specified (see Materials & Methods). The third and final type of input is what time points and populations should be saved, so `SOS` can later sample individuals from these and calculate different summary statistics based on those samples.

Once all three types of input are specified, `SOS` is used to simulate the epidemic with selection (Fig. 1B. step 1) as many times as the user wants. And importantly, it allows the user to subsequently sample from those simulations at one or more of the time points that were specified as part of the third input (Fig. 1B. step 2). Finally, `SOS` can then be used to calculate different selection scan statistics from the samples and thus to evaluate which ones can detect epidemic-driven selection given a particular sampling scheme (Fig. 1B. step 3). Currently, the selection statistics available within `SOS` include classic frequency-based methods such as the fixation site index (*F_ST_* ) and the joint site frequency spectrum (jSFS) as well as popular haplotype-based methods like iHS. Hence for example, the user can specify that they want to compare 100 samples from before versus 100 from after an epidemic with *F_ST_* . `SOS` will then perform both the sampling and a *F_ST_* selection scan and either visualise the selection scan results or summarise them into power estimates (Fig. 1C). Additionally, `SOS` can also provide the raw sampled output to make it possible for the user to perform additional statistical analyses of the sampled data if relevant.

Below we illustrate how `SOS` can be used in practice with two example epidemics. In both examples, for simplicity, we simulated data from two chromosomes, equal to human chromosomes 21 and 22 (a total of 68.9 Mb), and chose to consider the selected variant successfully detected if it was among the top three candidate variants or if one of the top three variants was in strong linkage disequilibrium (LD) with the selected variant (*r*^2^ *>* 0.8). Furthermore, for speed, we performed all our power analyses by first running 10 simulations for each epidemic scenario (i.e. combination of sample scheme, viability, sample size and starting allele frequency) each with a different selected locus followed by 100 subsamplings of each of those and then estimating power as the proportion of those 1000 simulation replicates per scenario. However, we note that the length and number of chromosomes simulated, the detection criteria, and the number of simulations performed can all be adjusted by the user.

### 2.2 Example of use: Rinderpest in the African Cape buffalo

The iconic Cape Buffalo went through a severe epidemic outbreak of rinderpest in the final decades of the 19^th^ century, which led to a high number of fatalities across the species range. Given the strong reduction in population size, approximately 90%, we were interested in exploring if this extreme epidemic outbreak drove positive selection on any single genetic locus. More specifically because we had 20 present day samples from Kruger National Park we wanted to use `SOS` to investigate if this was enough for a well powered study and if not whether, for example, adding more modern samples and/or some older samples from before the epidemic would lead to a useful study.

As mentioned earlier, to be able to use `SOS` three types of input are needed: (1) data that mimics the relevant populations(s) right before the epidemic-driven bottleneck, (2) a description of the epidemic-driven bottleneck and assumptions about the nature of the potential selection, and (3) a delineation of the populations and generations of interest for later analyses (Fig. 1). To answer our questions for the rinderpest epidemic in African buffalo, we therefore used `SOS` in the following way:

For the first input type, we simulated data that mimicked the population from which we had sampled (*n* = 20), Cape buffalo from Kruger National Park (KNP). To this end, we used a recently published PSMC [23] analysis when simulating using `SLiM` [22] the demographic history of KNP Cape buffalo while cross-checking our simulations with the present-day levels of heterozygosity of KNP Cape buffalo (see Materials and Methods).

For the second type of input, we compiled a combination of census data of KNP buffalo and previous work on rinderpest [24]. This provided us with an estimated size and length of the epidemic-driven bottleneck: about a 90% decrease in population which took place in the span of approximately 1 generation. Following this, we decided to perform a gradual recovery taking place across 15 generations post-epidemic to present-day. For the advantageous variant, we initially assumed an additive model of inheritance during the rinderpest epidemic and to get a broad overview, we explored all possible combinations of four different viabilities for the homozygous carriers (*V_AA_* = 1, 0.8, 0.5, and 0.3) and four different starting frequencies (*f_A_* = 0.1, 0.2, 0.3, and 0.4).

For the third input, we specified that we wanted to save the following time points of interest: the present plus the generations right before and right after the epidemic in case it turned out our available modern data were not enough for a well-powered study.

With these inputs we simulated selection during the epidemic using `SOS`. We then first investigated whether our 20 modern samples led to enough power by sampling 20 simulated samples from the present-day (saved by `SOS` during simulations) and applying the following statistics often used to detect selection using only present day samples, namely iHS, *π* and Tajima’s *D*. Since computing iHS is inefficient we chose only to focus on the simulations from the selection scenario with the strongest viability (*V_AA_*= 1). Unfortunately, even for this scenario our simulations and subsequent samplings show that there is insufficient power (*≤* 4.5%) to detect epidemic-driven selection using any of the applied statistics when looking solely at 20 present-day samples (Table S1).

Next, we tried to apply `SOS` to the same simulations, but using a much larger sample size (*n* = 1000) from the present day generation in order to get a sense of how many more samples we would need to obtain a well-powered study if we analyse only present-day samples. However, even at *n* = 1000 detection of the selected locus did not render sufficient power (*≤* 7% power) (Table S1).

Finally, we explored whether adding historical samples from before, and potentially also right after the epidemic, might help and if so, how many samples would then be needed. More specifically, we used `SOS` to estimate the power for two common selection statistics, *F_ST_*and jSFS, for the following two additional sampling schemes assuming a range of different sample sizes per group of samples being compared (*n* = 20, 50, 100, 200 and 500):

1. Before versus Present: Comparing individuals from before the rinderpest epidemic to the present-day population (i.e. 15 generations post-epidemic).
2. Before versus After: Comparing individuals from before the rinderpest epidemic to the individuals of the next generation.

We used `SOS` to estimate the power for each possible combination of simulated values of *f_A_* and *V_AA_*. We did this for both *F_ST_* and jSFS, however we will only describe the results for *F_ST_* here since the results for jSFS were in general similar to those of *F_ST_* and when different, *F_ST_* tended to perform better (Fig. S3).

Overall, the two comparative sampling schemes for the rinderpest epidemic in Cape buffalo yielded markedly increased detection power compared to performing selection scans solely on present-day samples (Fig. 2A). Moreover, they both led to some well-powered designs, especially when the advantage of carrying the advantageous allele was highest (*V_AA_* = 1): both Before versus After and Before versus Present had powers over 80% when using *F_ST_*for *n ≥* 50 samples from each of the time points (Fig. 2A). When the advantage is lower *V_AA_* = 0.8, 50 samples from each time is also enough to obtain a power of at least 80% for the Before versus After sampling scheme, but for the Before versus Present at least 100 samples from each time point are needed. For even lower viabilities, the number of samples required becomes highly dependant on the starting *f_A_* of the advantageous allele: the lower the starting *f_A_*, the lower the power. And for the lowest investigated starting *f_A_*, 0.1, the number of samples from each time point required to obtain a power of at least 80% are 200 and 500 for *V_AA_*= 0.5 and 0.3, respectively when using a Before versus After sampling scheme. Notably, *n* = 20 samples from each time point led to only modestly powered studies across all explored viabilities (e.g power of less than 59% using *F_ST_* ) for both sampling schemes.

**Figure 2:**
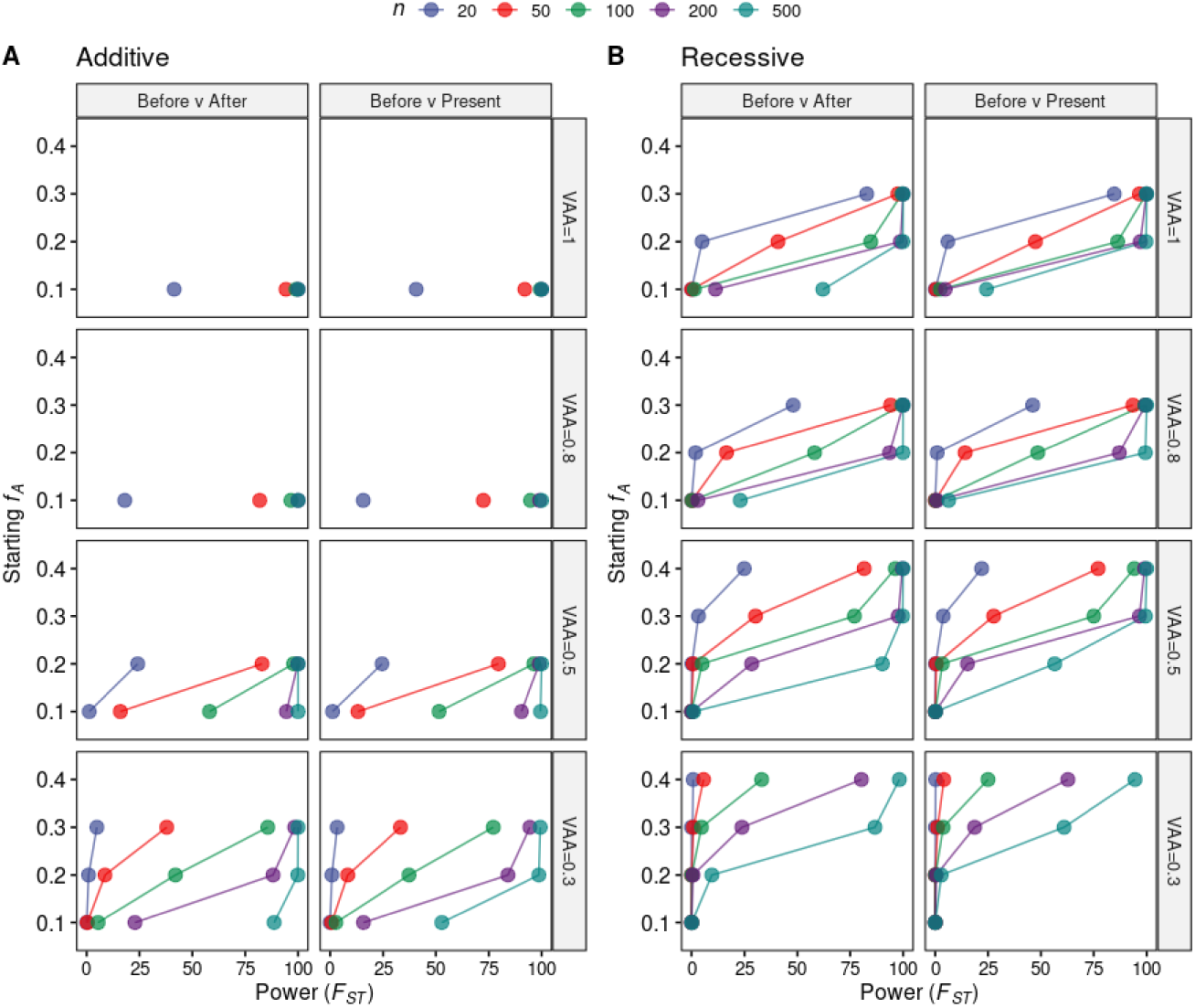
Simulation based estimates of detection power assuming additive and recessive selection in rinderpest in African Cape buffalo. Power to detect selection driven by the rinderpest epidemic with a 90% mortality estimated using `SOS` for different combinations of sampling schemes, initial allele frequencies of the advantageous allele (*f_A_*), viabilities of the homozygous carriers of the advantageous allele (*V_AA_*), and sample sizes for each grouping (*n*) (see Table S1 for exact values). Power was estimated as the percentage of simulation replicates, where the selected variant was detected from a total of 1000 performed simulation replicates per scenario with *F_ST_* for (**A**) additive and (**B**) recessive selection. Note that results for some initial allele frequencies are missing for some of the *V_AA_* values because those combinations are biologically unfeasible. For instance with an initial allele frequency of 0.4, then assuming Hardy Weinberg Equilibrium right before the outbreak, 16% of the population is expected to be homozygous carriers of the advantageous allele and that means 90% of the population cannot be killed if the viability of the homozygous carriers, *V_AA_*, is set to 1 (i.e. they all survive).

As stated in the beginning, we chose to focus on an additive model and thus all the results above are under that model. However, we also tried to perform the same simulations under a recessive model (Fig. 2B; Table S2). Interestingly, for the sampling scheme with only present-day samples there was actually one scenario (*V_AA_*= 1 and an initial *f_A_*= 0.3, where we obtained 63% power using iHS on *n* = 1000 present-day samples (Table S2). Though this could potentially be due to iHS being most powerful when the target allele has a frequency between 0.4 and 0.8 [25]. However, we note that all other tested scenarios under recessive selection rendered 0 or close to 0% detection power, suggesting that more than 1000 present-day samples would be needed to reach a well-powered study for this sampling scheme assuming a recessive inheritance model. For the comparative sampling schemes we note, not surprisingly, that more samples are required under recessive selection compared to under additive selection in all comparable investigated scenarios. Considering these results, if we wanted to design a study to look into the rinderpest epidemic in KNP buffalo, we would discard the possibility of solely using present-day data to detect epidemic-driven selection. The most optimal of the designs explored, power-wise, is having samples from before and right after the epidemic and using *F_ST_* . For this design we learned that, even under strong selection (*V_AA_*= 1), having at least 50 samples per comparative group is needed for a well-powered study (i.e. detection power *≥* 80%). Moreover, under weaker selection *V_AA_ ≤* 0.8 and smaller starting allele frequencies, even more samples would be necessary (Fig. 2; Table S1 and S2). Generally speaking, the largest *f_A_* feasible under any assumed advantage led to the highest power and therefore the most optimistic assumption with the difference in power being sometimes large: e.g. with *n* = 100 the power for *F_ST_* Before versus After when *f_A_* = 0.2 is

*≈* 44% of the power when *f_A_* = 0.3 assuming *V_AA_* = 0.3. Most promisingly, since modern samples are usually easier to obtain, our simulations showed only a modest difference in detection power between Before versus After and Before versus Present in either modes of selection.

### 2.3 Example of use: Black Death in Medieval Sweden

In our second example of how `SOS` can be used we explored an epidemic with different characteristics to those of rinderpest. Where rinderpest was a severe single generation epidemic in Cape buffalo, for our second example we explored a historical multigenerational epidemic that was comparatively less lethal, but with one of the largest death tolls in human history: the Black Death with a focus on Medieval Sweden.

The bacterium *Yersinia pestis* has been responsible for some of the most devastating pandemics in human history. Among the three major plague pandemics, the second plague is recognized as a turning point in European history where up to 60% of the European population is believed to have perished in less than a single generation [26]. Several studies have looked into whether Black Death could have posed a strong selection pressure upon variants that confer protection against *Y. pestis* in the European population [7, 9, 12], however clear, undisputed evidence of such selection has yet to be presented [13].

Much of the second plague pandemic research has concentrated on populations and regions where abundant multidisciplinary resources exist locally in relation to the Black Death [27–31], such as England, France and Italy. Comparatively less is known about the Scandinavian “Great Death”, as the second plague is referred to in contemporary Swedish.

Motivated by previous studies that have searched for putative plague-driven selection, we used `SOS` to explore when, and if, it would be possible to detect plague-driven selection in Medieval Sweden while mimicking the sampling schemes many related studies have used i.e. Before versus After [7–9, 12]. Additionally, we decided to explore a similar sampling scheme used in a few studies of epidemics in relation to the Black Death [8] but also used for exploring other acute epidemics, such as Ebola [6]: comparing the dead (non-survivors) to survivors. In principle, comparing those that died from the epidemic to those that survived should allow us to assess if a variant that putatively confers protection against *Y.pestis* is found in higher frequency among the survivors relative to the victims.

To explore these sampling schemes for Black Death in Medieval Sweden, we used the Gravel model [32] in `SLiM` to simulate the demographic history of a European-like population for our first `SOS` input (Fig. 1A). The second input type required compiling information about the plague in Sweden. We used historical records and accounts by various experts to estimate the size and length of the plague-driven bottlenecks in Sweden [26, 33, 34]: a 50% reduction in population size from 1350 CE until 1430 CE, caused by two outbreaks lasting a generation each and separated by one generation. We initially aimed to explore the same range of starting frequencies and viabilities for the potential advantageous variant as in our rinderpest example, but realised that the majority of lower viabilities led to no observable power. Therefore we only show results for all possible combinations of two different viabilities for the homozygous carriers (*V_AA_* = 1 and 0.8) and four different starting frequencies (*f_A_* = 0.1, 0.2, 0.3, and 0.4). We explored both an additive and a recessive model of inheritance selection for comparison purposes. For the third input of `SOS` we decided to save the following time points: the generation before the epidemic, during the second severe bottleneck, and right after the second bottleneck of plague.

With this input we used `SOS` to simulate and estimate power to detect selection (Fig. 1B. steps 1-3) with the selection statistic, *F_ST_* which is the most commonly used in previous studies. Fig. 3 displays the detection power for each scenario, mode of inheritance, and method estimated by `SOS`. Under an additive selection model, we note that in order to obtain a well-powered study (power *≥* 80%) using a Before versus After sampling scheme, at least 500 samples are needed in each grouping and that is under the assumption of complete protection when homozygous for the advantageous allele (*V_AA_* = 1.0) and a starting *f_A_*close to 0.3. Lower advantages of being homozygous for the advantageous allele (*V_AA_*= 0.8) led to virtually no detection power (*<* 5%) even at higher *f_A_* and larger sample sizes (Fig. 3 A) across all sampling schemes. On the other hand, when exploring the Dead versus Survivors sampling scheme, we observe that when homozygous individuals are completely protected by the advantageous allele (*V_AA_* = 1.0) and assuming an initial *f_A_* of 0.3 (Fig. 3), *n* = 100 per grouping is sufficient to render a well-powered study (power *>* 88% power across all comparative methods).

**Figure 3:**
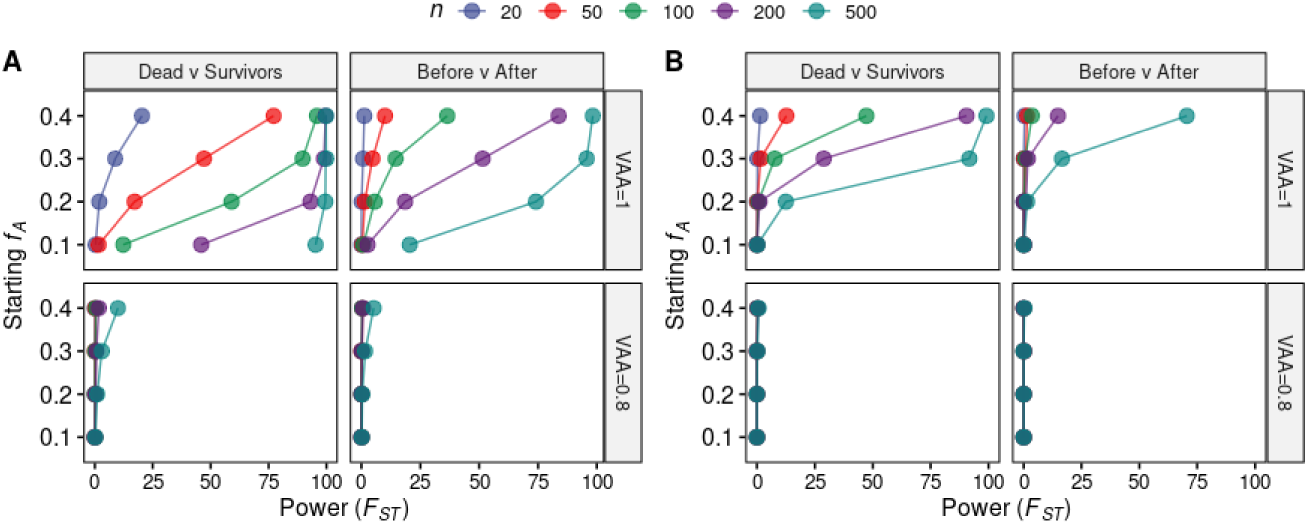
Simulation based estimates of detection power assuming additive or recessive selection in Black Death in Medieval Sweden. Different plague epidemic sampling schemes comparing initial allele frequencies (*f_A_*), sampling sizes for each grouping (*n*), and detection power using *F_ST_* for (**A**) additive and (**B**) recessive selection. Power values correspond to the percentage of simulations where the selected variant was detected from a total of 1000 performed simulation replicates per scenario. These simulations explored two different viability values for the homozygous advantageous allele (*V_AA_*) combined with four different initial allele frequencies.

Under recessive selection model, Before versus After as a sampling scheme gave rise to no well-powered designs (power *<* 80%), suggesting only higher initial allele frequencies could render a study well-powered assuming recessive selection and with a substantial amount of samples (*n >* 500) (Fig. 3B; Table S3). The Dead versus Survivors sampling scheme rendered well-powered designs but requires more samples compared to an additive model. Particularly we note that more than 500 samples per grouping is necessary to reach reasonable power if the initial *f_A_*is 0.1 (Fig. 3B; Table S3).

Overall, simulations of the Black Death in Medieval Sweden suggests that only a *V_AA_*= 1 renders any power to detect epidemic-driven selection with the explored sample sizes. Hence, unless selection was extremely strong, the amount of samples needed for a well-powered study is very large, especially given the sampling schemes explored are based on historical samples. Notably, the optimal sampling scheme for this epidemic was Dead versus Survivors whereas many more samples were necessary to detect the protective variant for Before versus After (Fig. 3).

### 2.4 Example of use: Gaining further insights

As a third example of how `SOS` can be used, we used some of the additional output that `SOS` can provide to gain some further insights into why detection power varies extensively between the different scenarios and methods for our two example epidemics.

#### Allele frequency shifts

First, from the results for the two examples of epidemics we just presented it was clear that, as expected, sample size and viability of the homozygous carriers matter substantially for power. But it was also clear that other factors, like starting *f_A_* and choice of sampling scheme also play a role. To get a better understanding of why, we first investigated the raw data from the different samples.

In particular, for all the scenarios with *V_AA_*=1, we calculated the mean difference in the frequency of the advantageous allele (Δ*f_A_*) due to additive selection for the different sampling schemes in each epidemic and plotted this against the estimated power for *F_ST_* -based scans (Fig. 4A and B). Unsurprisingly, we see a clear correlation: the larger the allele frequency difference (Δ*f_A_*), the higher the power. However, focusing only on the simulations from the plague epidemic (Fig. 4), we also observe that the increase in power can, to some extent, be explained by the fact that higher starting allele frequencies result in larger differences in the allele frequency of the advantageous allele, *f_A_* due to selection.

**Figure 4:**
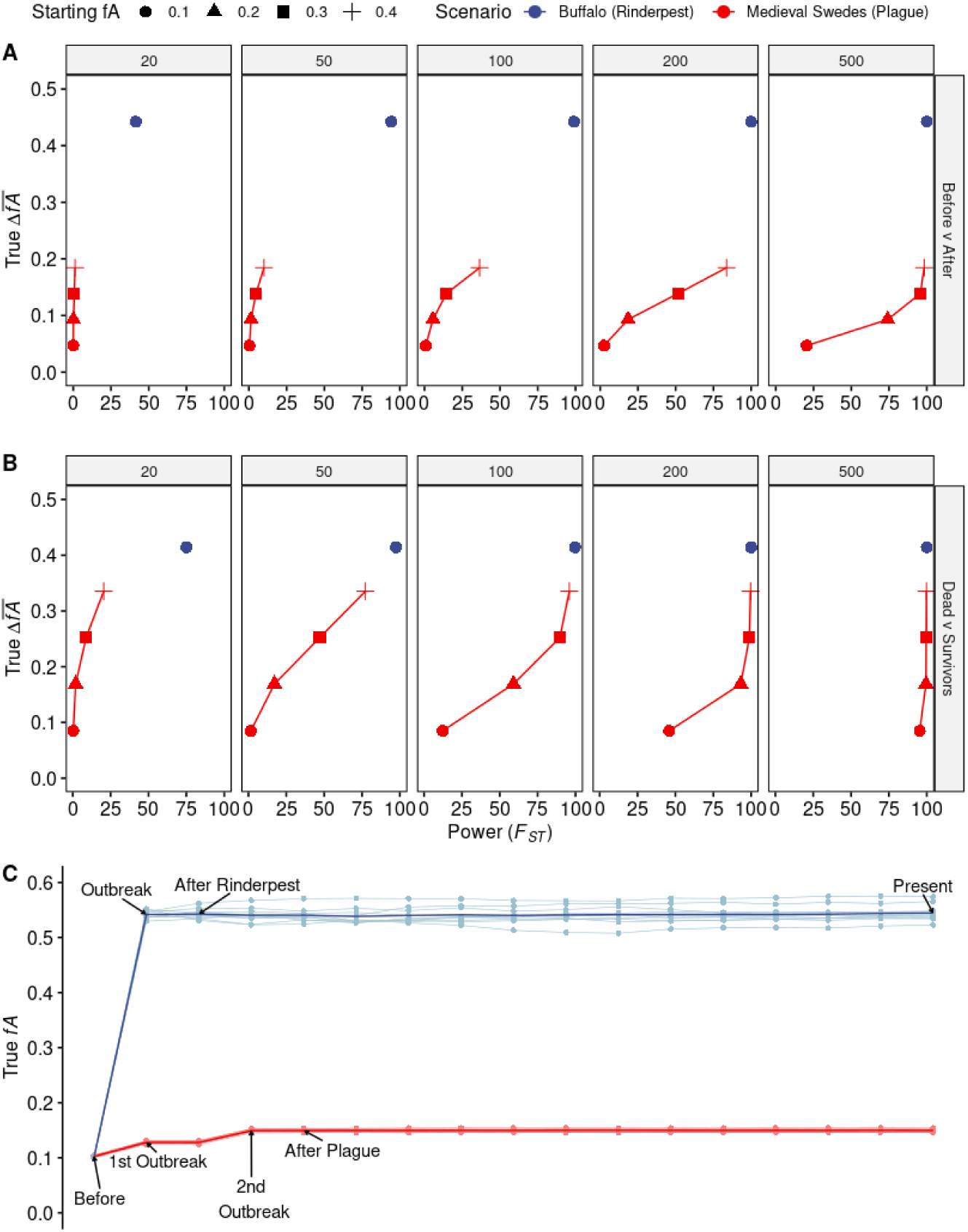
Allele frequency differences observed for the rinderpest and plague epidemics assuming. *V_AA_* = 1. Differences in allele frequency, Δ*f_A_*, are plotted against detection power using *F_ST_* for sampling schemes (A) Before versus After and (B) Dead versus Survivors in all scenarios where being homozygous for the advantageous allele gives an individual the highest advantage (*V_AA_* = 1). In the results the two epidemics are separated by color, initial allele frequency by shape, and each panel is divided into sample size used (*n* = 20, 50, 100, 200, 500). (C) True *f_A_* trajectories across time in the 10 simulations for rinderpest in Buffalo (light blue) and for plague in Medieval Swedes (light red) for a starting *f_A_* of 0.1; mean *f_A_* is coloured by epidemic (dark blue and dark red).

**Figure 5:**
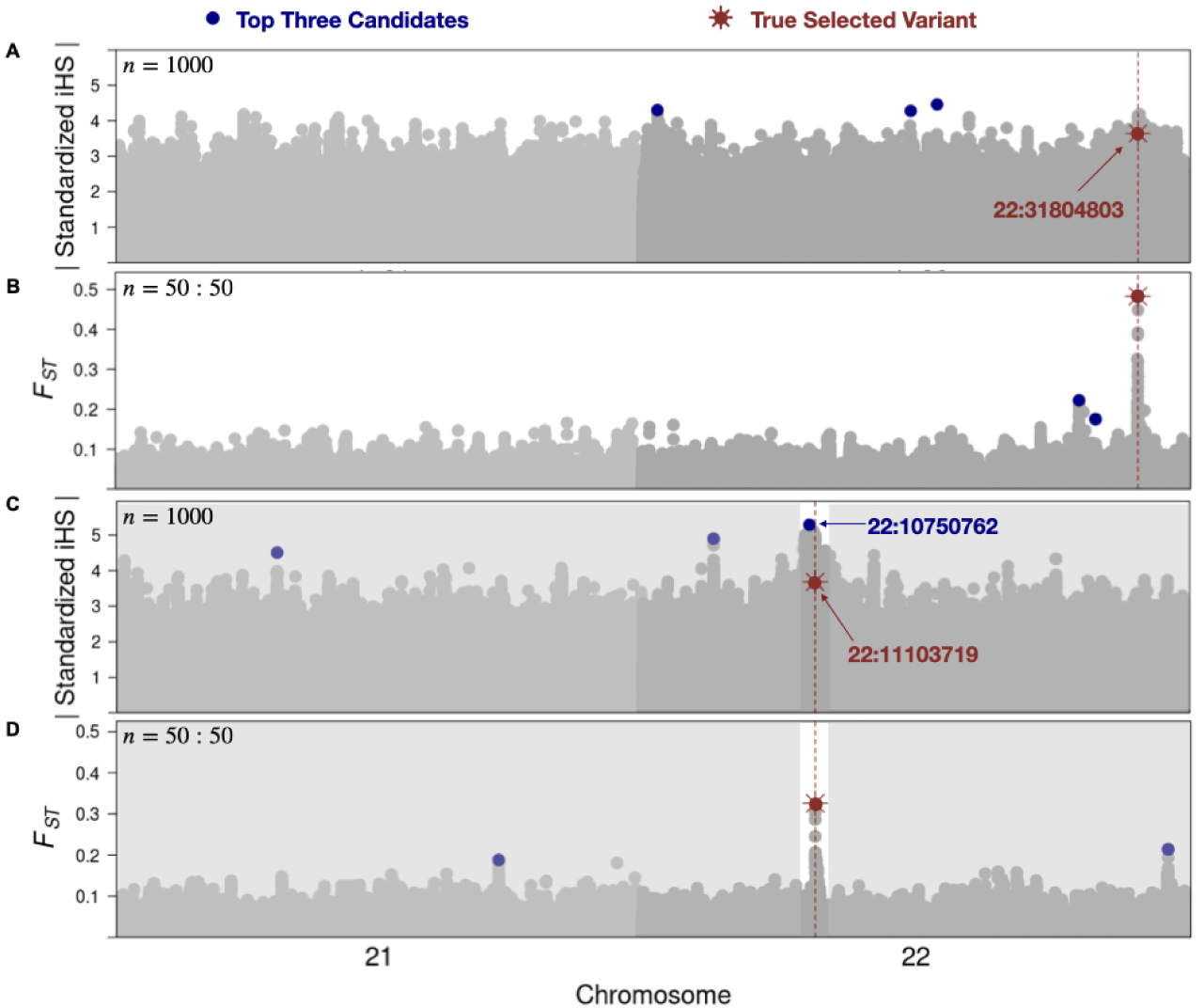
iHS and *F_ST_* graphical output by `SOS` comparing results for a rinderpest simulation under additive selection, with *V_AA_*=1 and *f_A_* = 0.1. iHS was performed with *n* = 1000 for the Present while *F_ST_* was performed on Before versus Present with *n* = 50 for each time point. (A) An example iHS result that shows no discsernible peak or nearby candidate while (B) using *F_ST_* shows clear detection of the selected variant. (C) An example iHS result that shows a broad peak where the top candidate detected was more than 350 Kb away (blue point) from the selected variant (red star) and with *r*^2^ *<* 0.1. (D) Corresponding *F_ST_* result for the broad peak showing better precision than iHS.

Furthermore, when we compare the plots for Before versus After and Dead versus Survivors, it reveals that the increased power in the latter sampling scheme can be attributed to the larger allele frequency differences it results in.

Finally, it is worth noting the differences in Δ*f_A_* for the plague epidemic and rinderpest epidemics, specifically for the case where the starting *f_A_* is the same (red vs blue circles in the plots). This clearly explains much of the big difference in power between the two epidemics: again, the larger the Δ*f_A_*, the higher power. And if one takes a look at the models used by `SOS` for rinderpest and plague the difference in Δ*f_A_* can be explained by the fact that the bottleneck sizes are very different: the starting frequency *f_A_* and the viability for the homozygous carriers of the advantageous variants *V_AA_* are set to the same, so to achieve different bottleneck sizes the viabilities of *V_Aa_*and *V_aa_*differ greatly. Specifically, the larger the bottleneck, the smaller the viability has to be for *V_Aa_* and especially *V_aa_*, which means the overall selection pressure gets stronger, which in turn leads to a larger Δ*f_A_*. Importantly, this overall trend is further supported by the true *f_A_* trajectory observed for each of the two epidemics simulated assuming *V_AA_* = 1 and *f_A_* = 0.1 (Fig. 4C). Here we see that the shift in allele frequency is much smaller in the simulations of the plague, which explains why more samples are required to detect selection in the plague epidemic compared to rinderpest under the same assumptions about *V_AA_*and starting *f_A_*.

#### Detection Criteria

The majority of the work presented here has focused on comparative sampling schemes. This is due to the little to no power we observed when performing single-population neutrality statistics (Table S1).

As a final example of how `SOS` can be used we tried to use `SOS`’s graphical outputs to evaluate why iHS consistently failed to detect the variant under selection, especially when we uphold the same criteria as we have for the comparative selection scans to consider a selected variant as detected (i.e. the 3 variants with the highest value of the relevant selection statistic being among or in high LD with the selected variant).

Specifically, we examined some selection scan plots from simulations with *n* = 1000 that `SOS` can output. Notably, in some simulations there is no signal at all surrounding the variant under selection, even when there was a clearly detectable signal using *F_ST_* based on samples sizes of only *n* = 50 per time point (Fig. 5A and B). However, we also found examples of simulations where the selected variant is within a broad peak that spans more than 0.35 Mb but where it was not considered detected according to our detection criteria (Fig. 5C). This implies that there are cases where we were unable to detect the selection using our detection criteria, but where it is possible to choose a less stringent criteria and detect the general region of interest. Naturally, the optimal choice of detection criteria depends on the aims of the user, but if detecting the region of interest without capturing a signal in the advantageous variant is a reasonable outcome`SOS` could be used to design a criteria that would increase the power of iHS applied to modern samples.

Inspired by this, we also further explored the effect of using a different detection criteria for *F_ST_* based detection. Specifically, we tried to compare the power obtained with the LD-based detection criteria used so far with the power obtained using a genomic distance based detection criteria, where the advantageous variant is considered detected if it falls within a 1Mb window of one of the top three variants (500 Kb on each side of the top variants). We observed that this new criteria in general led to increased power. Across all scenarios explored the increase in power ranged 0 *−* 27.4% in rinderpest (Fig. S5) and 0.1 *−* 16.8% in plague (Fig. S6). However we note that the median increase is only 6.2% and 5.3%, respectively, and only in a very few scenarios the increase made a difference in the conclusion of whether the study was considered underpowered (i.e. *<* 80% power) (Fig. S7-8).

## 3 Discussion

Here we have presented `SOS`, a simulation-based framework that allows users to explore whether epidemic-driven selection can be detected for different epidemic scenarios, sampling schemes, and sample sizes. Furthermore, we have showcased, using two example epidemics, how our framework can be used to inform users about when and, if, they have reasonable power to detect epidemic-driven selection.

Besides illustrating how `SOS` can be used, the results from our two example epidemics highlight several important points. First, our results reveal that power is highly dependent on the size of the bottleneck and therefore varies greatly between different epidemics. Despite the shared parameters the two epidemics were simulated with (i.e. *f_A_*, *V_AA_*, mode of inheritance), the different bottleneck sizes cause marked dissimilarity in allele frequency changes, and therefore in power (Fig. 4).

Secondly, our simulations also show that unless the bottleneck size is extreme, like for the case of the rinderpest epidemic in Cape buffalo, a substantial number of samples appear to be necessary to obtain reasonable power (*≥* 80%). And even in an extreme case, like rinderpest, we note that we were unable to detect the epidemic-driven selection using only 20 present-day samples. While a number of previous studies have attempted to detect signatures of a rinderpest-driven bottleneck in African buffalo [23, 24, 35–38], few have specifically sought out signatures of selection driven by the epidemic [2]. Notably, the previous studies have shown that neither genomewide heterozygosity nor site frequency spectra were altered by the rinderpest bottleneck, which may be what has discouraged a focus on selection. However, we too find that despite simulated high death-tolls, there is no marked difference when using standard measures of diversity or population differentiation. Importantly, despite this, with sufficient samples (*n ≥* 50) and a comparative sample scheme, we can still detect signatures of rinderpest-driven selection. This suggests that a study with focus on selection driven by rinderpest may be worth performing.

Thirdly, not all the explored comparative sampling schemes lead to similar power. This is especially highlighted in our results for the plague epidemic in Medieval Sweden where sampling scheme mattered considerably. Specifically, Dead versus Survivors rendered markedly better power than both Before versus After or Before versus Present. On the other hand, the difference between Before versus After and Before versus Present was less clear in the case where we made that comparison. This is encouraging since samples from the present are often easier to get access to than samples from right after the plague. However, we caution that this conclusion of course depends on if and how the population’s size recovers after the epidemic.

Fourthly, we also observe that one assumption about mode of inheritance matters greatly. For example, assuming additive selection would lead to over optimistic power estimates (Fig. 2; Fig. 3) if the true inheritance model is recessive and therefore misleading in how many samples would be necessary to have a well-powered study.

Finally, it is also important to note that we have simulated plague scenarios where a putative monogenic variant would be exceptionally protective in the population (i.e. *V_AA_* = 0.8, 1). Nonetheless, there are several combinations of viabilities and initial *f_A_* where little to no power could be obtained (Fig. 3). If achieving sufficient power to detect a protective monogenic variant requires a large sample size, it is conceivable that detecting polygenic epidemic-driven selection would require a substantially larger number of samples. This is an especially relevant consideration given that the genomic basis for host susceptibility to infectious diseases is largely polygenic [39, 40].

Together this suggests that running `SOS` before carrying out a study can be very useful if one wants get an idea of whether the samples they have are sufficient to perform a well-powered study. It also suggests that several previous studies may have been underpowered [13]. In fact, the largest plague study to date, in regards to pre- versus post-, has been with 206 individuals in total: 143 from London (38 pre; 42 during; 63 post) and 63 from Denmark (29 pre; 34 post) (Table S9). Although to fully conclude if such studies are underpowered, simulations specific to those exact epidemics would be needed. Importantly, if those studies were indeed underpowered, the fact that they did not find any signatures of epidemic-driven selection does not necessarily rule out the possibility that such selection took place. Hence larger future studies of the same epidemics could still be interesting to conduct.

It should be noted that `SOS`, and the results presented here, do have some limitations. To begin with, `SOS` requires different curated inputs (Fig. 1A) making it non-trivial to use. This includes having to know specific information regarding the epidemic one wishes to simulate. Moreover, we have shown with our results how the criteria for finding a selected variant plays a crucial role in detection power (Fig. 5 and Fig. S7-8). Unsurprisingly, we observe that by using alternative criteria, such as physical distance over LD, one’s power of detection increases. However, it of course comes at the cost that the causal variant may be difficult to find. Importantly, all our presented results were based on a naive outlier approach, mainly chosen because it is the most commonly used approach in similar studies [6–9]. However, being an outlier is evidently not enough to determine whether a variant has been under selection, and therefore it is not necessarily the most optimal approach. A potential alternative, especially in the case of the Dead versus Survivor sampling scheme, is to use *p*-values from per SNP association tests, which is also implemented in `SOS`.

Another limitation is that currently `SOS`’s simulations builds on the assumption that everyone in the population was exposed to the disease-inducing pathogen. Simulating parts of the population that are not exposed would be a more realistic way of modelling an epidemic and a relevant consideration for power. It is also worth noting that while we explore recessive and additive models of selection, other modes of inheritance can be appropriate if there is already known information about the host’s susceptibility to the pathogen. Furthermore, our results have been generated under the assumption of perfect data. This means that we have simulated perfect genotypes. If this is a concern it should be possible to add a layer of errors with existing tools like NGSNGS [41], so that the simulated data better mimics the actual data a user has available, including low depth, DNA damage, contamination, etc. While this has not been implemented in this study it is possible to do so using the raw outputs produced by `SOS` and this could help prevent overly optimistic power estimates. Finally, in our simulations we have assumed that we have accurate information about who has died of the epidemic and who has survived. This may of course not be the case. For this reason `SOS` has a built-in option called “mixed cemetery”, which allows users to specify a fraction of the individuals that perished from the epidemic and the individuals that survived to be swapped, thereby mimicking the uncertainty in determining cause of death in a burial site. This could be relevant to use if there is a substantial concern that the determination of the cause of death is not accurate.

From a broader perspective, we note that `SOS` can also be used to explore power for other critical events with significant death-tolls over a short time, besides epidemics. Recent short-term selection is of strong relevance to a wider community interested in understanding the impact anthropogenic disturbances has had on several species. Some of these phenomena include tusklessness in elephants due to drastic ivory pouching [42], pesticide-driven population declines in honeybees [43], and the cattle anti-inflammatory drug threat to white-rumped vultures [44, 45]. Users interested in similar episodic pressures would also be able to use `SOS` to design a study with appropriate power.

In conclusion, while `SOS` needs to be used with care and can be further improved, the examples in this paper clearly show that it can be highly beneficial in designing well-powered studies. Therefore, we believe and hope that `SOS` provides a valuable step in the long-term journey towards understanding the extent that epidemic-driven selection has affected present-day populations.

## 4 Materials and Methods

### 4.1 Description of the selection models used

#### 4.1.1 Notation

In our epidemic-driven selection model we work with viabilities and use the following notation:

We are focusing on a locus with three genotypes *aa* (WT), *Aa* (heterozygous carrier of an advantageous mutation) and *AA* (homozygous carrier of an advantageous mutation). In this locus, we denote the number of individuals in the three genotype categories in generation 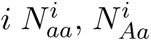 and 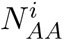 . We also denote the viabilities of the three genotypes *V_aa,_ V_Aa_* and *V_AA_*. Note that these translate into relative fitnesses with the following selection coefficients:

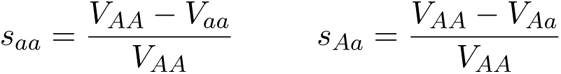

using the the following relative fitness model:

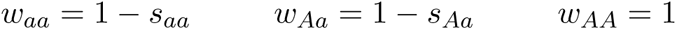

#### 4.1.2 Recessive, additive and dominant selection

With this notation then if we have a recessive model with *s_aa_* = *s_Aa_* we get the viabilities *V_aa_*= *V_Aa_* as this gives us

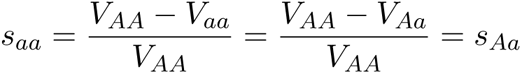

Similarly, if we have an additive model with *s_aa_* = 2*s_Aa_* we get that *V_AA_−V_aa_*

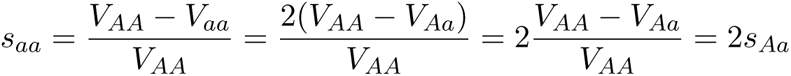

In other words in an additive model the difference in viabilities of the genotypes *aa* and *AA* (*V_AA_ − V_aa_*) has to be twice as high as the difference in viabilities of the genotypes *Aa* and *AA* (*V_AA_ − V_Aa_*).

Finally, if we have a dominant model with *s_Aa_* = 0 (and thus *w_Aa_* = 1 *− s_Aa_* = 1 = *w_AA_*) we get the viabilities *V_Aa_*= *V_AA_* as this gives us

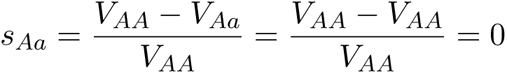

#### 4.1.3 Calculating viabilities that give rise to a specified bottleneck size

Now assume we need to reduce the population size to *X* via viabilities while taking into account the genotype counts. I.e. so

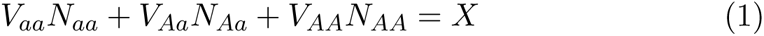

where *X* is a specific populations size.

Under **recessive selection** we want to find *V_aa_* and *V_Aa_* and we know that *V_aa_*= *V_Aa_* and therefore we can insert that into equation (1) and solve for *V_Aa_*:

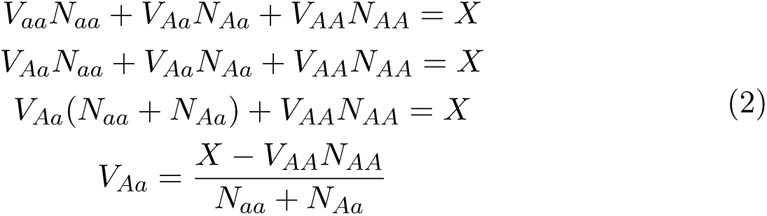

which means we have an expression for *V_Aa_* (and thus *V_aa_*) if we know *X*, *N_aa_*, *N_Aa_*, *N_AA_*, and *V_AA_*.

For an **additive model** we know per definition that *V_AA_*-*V_aa_* = 2(*V_AA_*-*V_Aa_*). This means that

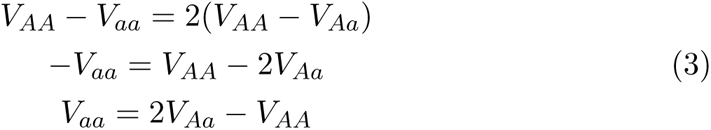

which we can then plug into equation (1):

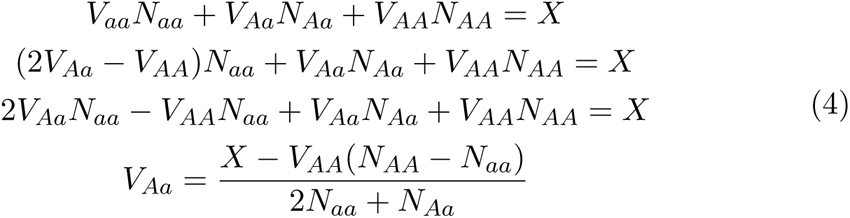

Similar to recessive selection, we know that in a **dominant model**, *V_Aa_* = *V_AA_* as per definition and thus we can also use this in equation (1) to solve for *V_aa_*:

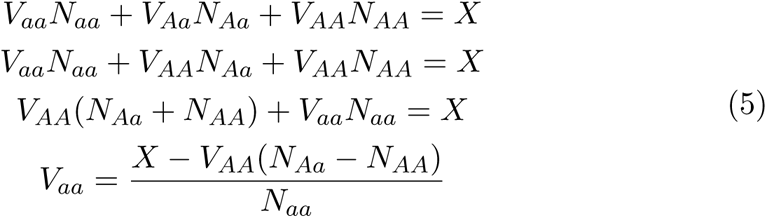

### 4.2 Simulations

All simulations were performed in `SLiM` [22] with tree-sequence recording. We first simulated the demographic history of each organism under a Wright-Fisher model as described in below for Cape Buffalo and Medieval Swedes. The recorded tree-sequence was then used as input into `SLiM` to simulate the epidemic bottleneck under a non Wright-Fisher model as detailed below for rinderpest and plague. For the epidemic phase of the simulation, we maintained non-overlapping generations but simulated viabilities (absolute fitnesses in `SLiM`) parameterised by the size of the epidemic bottleneck, the assumed selection model and strength (see above Methods 4.1), and the initial frequency of the target variant. We simulated two chromosomes in `SLiM` with a total length of 6.89 Mb. Following the completion of each epidemic forward-simulation, we recapitated the population using `msprime` [46] with the respective ancestral population size, recombination rate, and mutation rate for each organism. For each epidemic scenario—rinderpest and plague—we performed 10 replicate simulations. Each of these simulations were subsampled 100 times, leading to a total of 1000 replicates for each combination of sampling scheme and parameters (*f_A_*, *V_AA_*, *n*).

We performed the following genome-wide selection scans and neutrality statistics on the simulation replicates: *F_ST_* , jSFS, iHS, *π*, and Tajima’s *D*.

#### 4.2.1 African Cape buffalo and rinderpest

##### Demography

We used the demographic history (Fig. S1) estimated for the KNP Cape Buffalo by Quinn et al. [23]. Specifically we used the estimates for population sizes for the first 40,000 generations, assuming a generation time of 7.5 years based on previous studies of Cape buffalo [24]. We initiated simulations with a population size of 100,000 diploid individuals, a mutation rate of 1.5*×*10*^−^*^8^, and a uniform recombination rate of 1 *×* 10*^−^*^8^. Specifics on the population splits, expansions, and bottlenecks can be seen in Fig. S1 and in the Github repository.

##### Epidemic

We used population sizes estimated from census data on Cape buffalo published in a previous study [24]. In particular we simulated the epidemic to last a single generation (generation 40,002) and gradually recovered the population to present-day sizes in 15 generations. We saved the population data in a `tree sequence` for the following time points: Before (40,000), During (40,002), After (40,003), and the Present (40,017). For downstream analyses we further divided the During time point into Dead and Survivors. Specifics of the simulation set up can be seen in Fig. S2 and in the Github repository.

#### 4.2.2 Medieval Sweden and *Y.pestis*

##### Demography

The demographic history for Medieval Sweden was simulated using the Gravel model [32] as provided by `SLiM` recipes. We used a variable recombination rate using the HapMap phase II recombination map for hg19 and a constant mutation rate of 2.36 *×* 10*^−^*^8^. We took the European-like population in the model and exponentially grew it using the same expansion rate as in the original Gravel model until reaching the respective population size for Medieval Sweden before the epidemic (700,000).

##### Epidemic

From historical records and relevant literature [33, 34], we simulated the plague epidemic as multigenerational taking place in two different generations which, together lead to a 50% reduction in the population when compared to the generation before the epidemic (58796). A general depiction of the plague model can be seen below in Fig. S4. Specifics for the plague epidemic simulation set-up can be found in Table S8.

#### 4.2.3 Pairwise population statistics: F**_ST_** and jSFS

Across all comparative sampling schemes we calculated Hudson’s *F_ST_* [47] and jSFS using `scikit-allel` [48]. We calculated these two statistics per SNP. If the target variant itself or if a site was in strong LD (e.g. *r*^2^ *>*= 0.8) with the target variant was among the top three candidates, the variant was then considered detected for that respective method in that respective simulation replicate. Each candidate had to be at least 1 Mb away from each other. We performed in-house tests (data not shown) to choose which number of top candidates was appropriate for the length of the simulated genome when considering a variant as a true detected outlier; i.e. too many top candidates can lead to detecting the selected variant by chance.

Specifically for jSFS, in addition to the aforementioned conditions, we considered an outlier a top candidate by using a two-dimensional kernel density estimation [49] whereby the least dense outliers were considered.

#### 4.2.4 Single population statistics: iHS, ***π***, Tajima’s ***D***

Single-population statistics (e.g. iHS, *π*, Tajima’s *D*) were performed on samplings from the Present generation. We computed iHS using `rehh` [50] with a minor allele frequency filter of 0.05 (`min maf=0.05`) and allele frequency bin size of 0.01 (`freqbin=0.01`). Notably we performed this selection scan on only a few scenarios (rinderpest; *n* = 20 and *n* = 1000) because of how long and computationally demanding it was across 1000 replicates (i.e. 524.056 hrs using 75 cores).

We computed Tajima’s *D* and *π* with a 10 Kb sliding window and a 2 Kb step using `scikit-allel`.

### 4.3 Code Availability Statement

`SOS` can be installed using conda or mamba. The code developed for the `SimOutbreakSelection` (`SOS`) framework, simulating data, and analysis presented here can be found at the GitHub repository: https://github.com/santaci/SimOutbreakSelection

## 4.4 Acknowledgements

Many thanks to Professor Janken Myrdal for providing historical context and population size estimates of Medieval Sweden during the Black Death. Thanks to the CPH PopGen group for useful discussions and feedback.

C.G.S. and I.M. were funded by the European Research Council (ERC-2018-STG-804679) awarded to I.M.

