## Supplementary Material for "Detecting epidemic-driven selection: a simulation-based tool to optimize sampling design and analysis strategies"

### 1 Supplementary Tables: Captions

Table S1: **Rinderpest Additive Selection.** Different rinderpest epidemic sampling schemes comparing initial allele frequencies ($f_A$), sampling sizes ($n$), and methods of detection. Values correspond to the percentage (%) of simulations where the selected variant was detected from a total of 1000 performed simulations per scenario. These simulations explored the viability for the homozygous advantageous allele ($V_{AA}$) set across different initial allele frequencies. Selection coefficients are based on $V_{AA}$ of 1, 0.8, 0.5, 0.3 set to $f_A$ of 0.1, 0.2, 0.3, where possible. Dashed columns suggest biologically unfeasible fitness and initial allele frequency combination.

Table S2: **Rinderpest Recessive Selection.** Different rinderpest epidemic sampling schemes comparing initial allele frequencies ($f_A$), sampling sizes ($n$), and methods of detection. Values correspond to the percentage (%)of simulations where the selected variant was detected from a total of 1000 performed simulations per scenario. These simulations explored the viability for the homozygous advantageous allele ($V_{AA}$) set across different initial allele frequencies. $f_A$ of 0.1, 0.2, 0.3 were set to $V_{AA}$ of 1, 0.5, 0.3 where possible. Selection coefficients ($s_{Aa}$) are based on $V_{AA}$ of 1, 0.5, 0.3 set to $f_A$ of 0.1, 0.2, 0.3, where possible.

Table S3: **Plague Recessive Selection.** Different plague epidemic sampling schemes comparing initial allele frequencies ($f_A$), sampling sizes ($n$), and methods of detection. Values correspond to the percentage (%) of simulations where the selected variant was detected from a total of 1000 performed simulations per scenario. These simulations explored the viability for the homozygous advantageous allele ($V_{AA}$) set across different initial allele frequencies. Selection coefficients ($s_{Aa}$) are based on $V_{AA}$ of 1 and 0.8 set to $f_A$ of 0.1, 0.2, 0.3.

Table S4: **Rinderpest Frequencies for Additive Selection.**

Table S5: **Rinderpest Frequencies for Recessive Selection.**

Table S6: **Plague Frequencies for Additive Selection.**

Table S7: **Plague Frequencies for Recessive Selection.**

Table S8: **Swedish plague simulation set-up and corresponding times.**

Table S9: **List of epidemic-selection studies and their respective study designs.**

### 2 Supplementary Figures

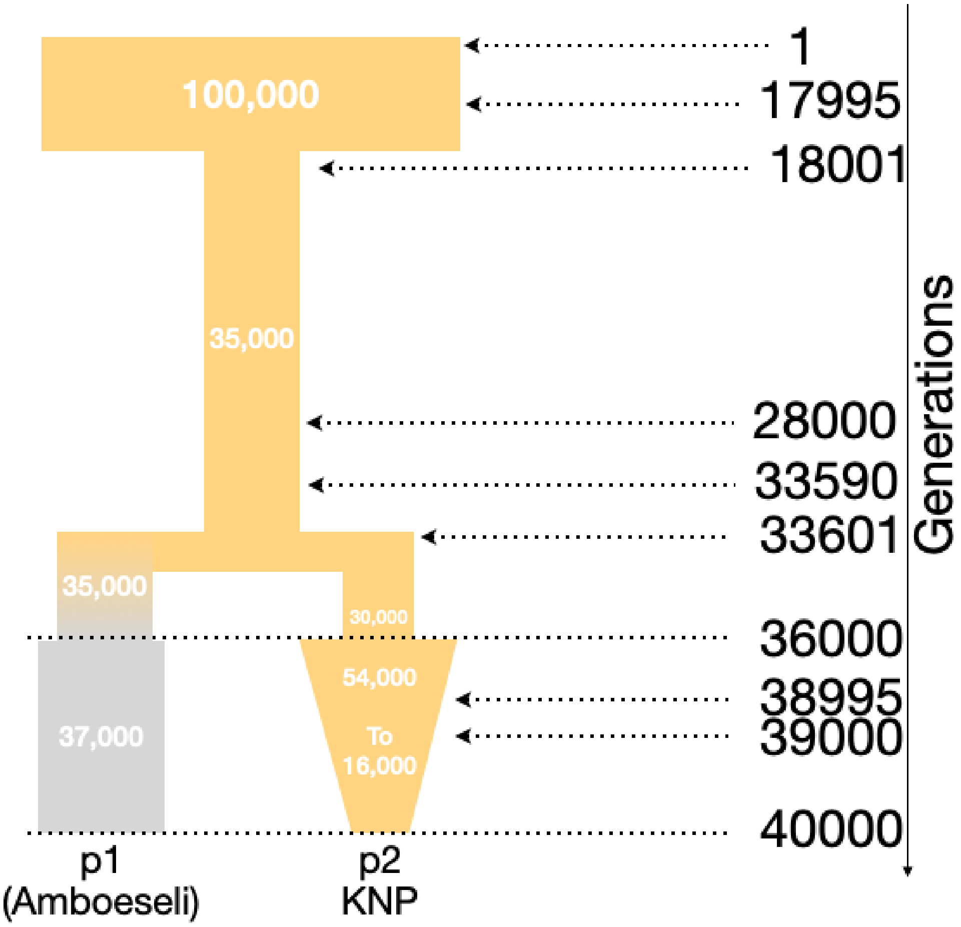


Figure S1: Wright-Fisher simulation set up for African Cape buffalo using `SLiM`. White numbers refer to population sizes. Y axis on the right refers to generation time going from the deep past to the recent past (top to bottom).

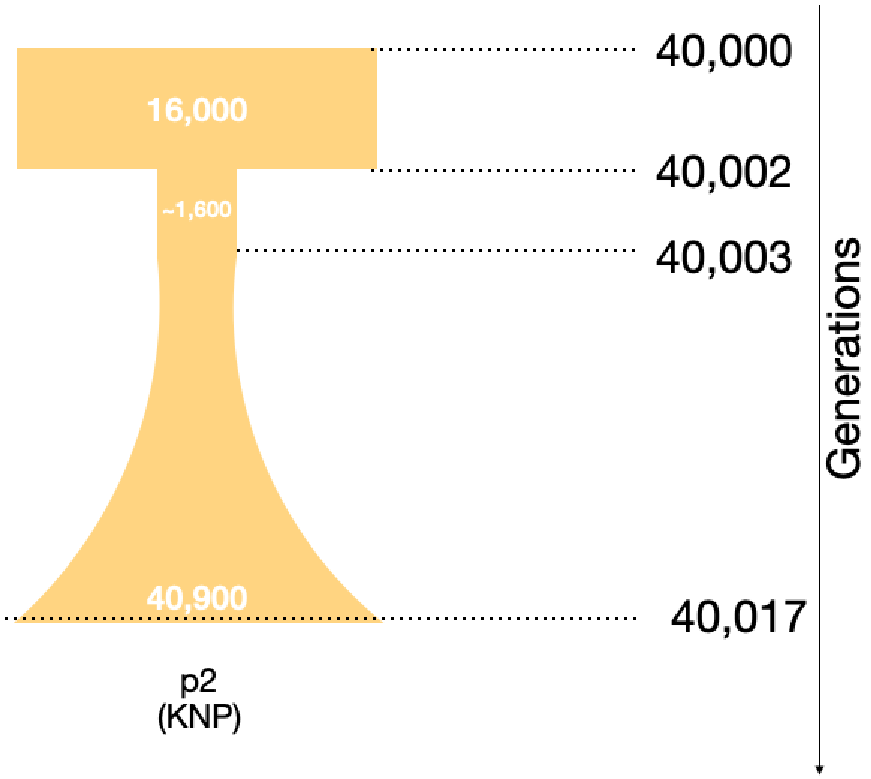


Figure S2: non Wright-Fisher simulation set up for rinderpest in African Cape buffalo using `SLiM`. White numbers refer to population sizes. Y axis on the right refers to generation time going from the recent past to the present (top to bottom).

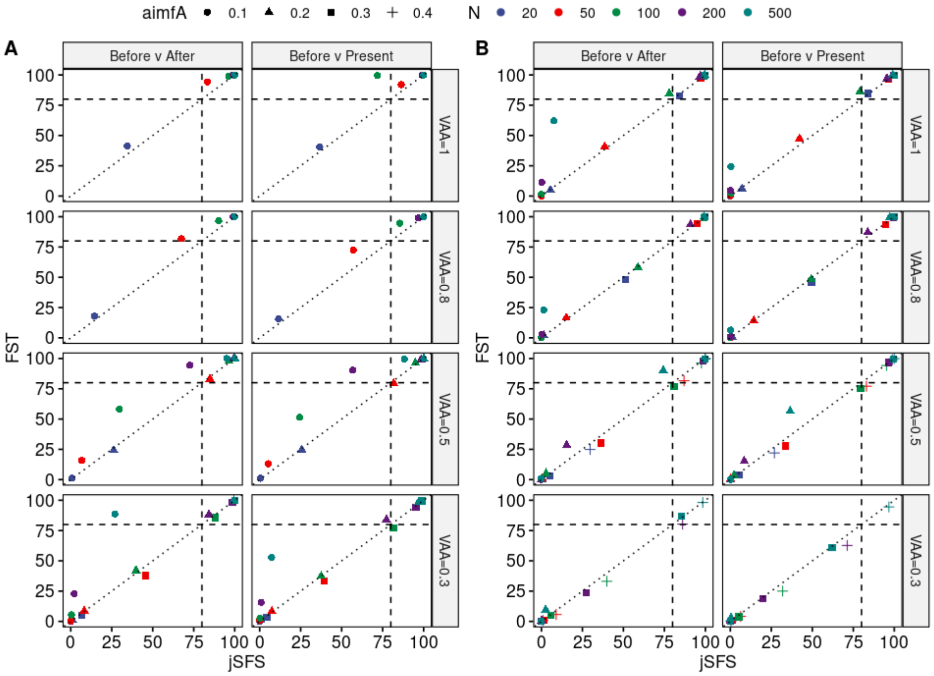


Figure S3: Scatterplot comparing detection power using jSFS and $F_{ST}$ in rinderpest in Cape Buffalo simulations under an **A.** additive and **B.** recessive model.

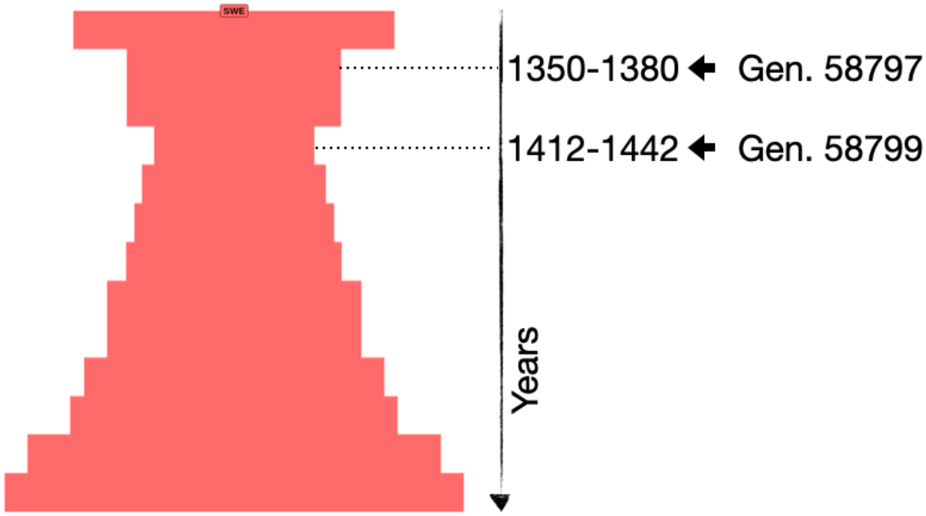


Figure S4: non Wright-Fisher simulation set up for plague in Medieval Sweden using `SLiM`. Y axis on the right refers to years and corresponding generations going from before the epidemic to the present (top to bottom).

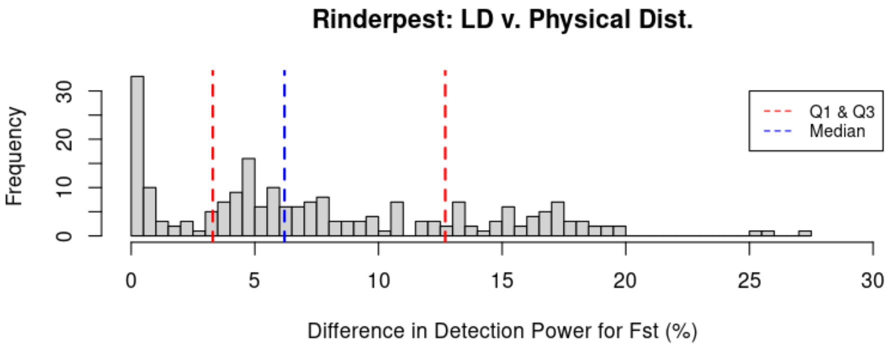


Figure S5: Difference in power of detection using $F_{ST}$ based on LD versus genomic distance—1 Mb (within 500 Kb on each side of the selected variant)—in rinderpest simulations for all pooled scenarios (additive and recessive).

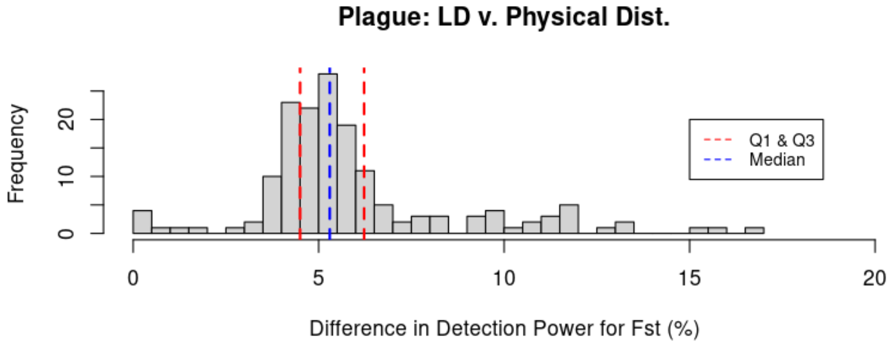


Figure S6: Difference in power of detection using $F_{ST}$ based on LD versus genomic distance—1 Mb (within 500 Kb on each side of the selected variant)—in plague simulations for all pooled scenarios (additive and recessive).

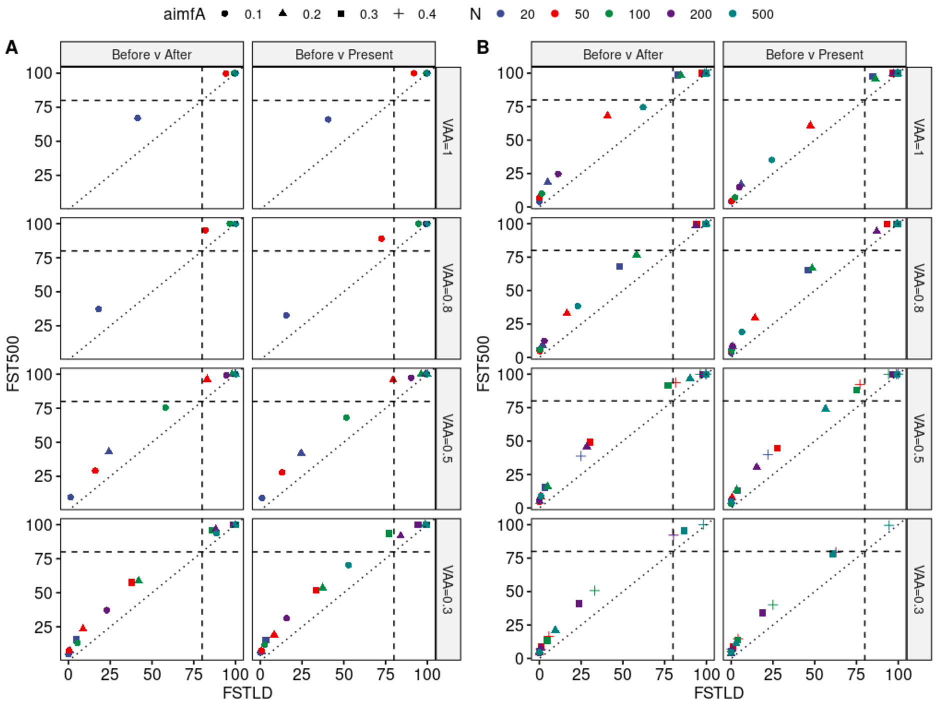


Figure S7: Scatterplots comparing power of detection using $F_{ST}$ based on LD versus genomic distance—1 Mb (within 500 Kb on each side of the selected variant)—in rinderpest simulations for **A.** Additive and **B.** Recessive models. Vertical and horizontal dashed lined represent 80% power threshold.

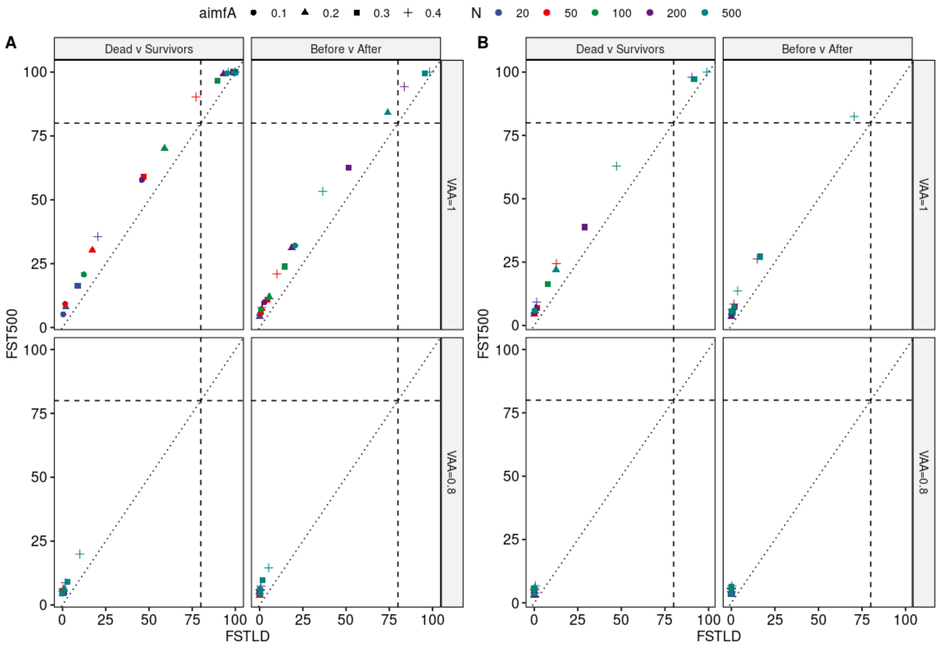


Figure S8: Scatterplots comparing power of detection genomic $F_{ST}$ based on LD versus physical distance—1 Mb (within 500 Kb on each side of the selected variant)—in plague simulations for **A.** Additive and **B.** Recessive. Vertical and horizontal dashed lined represent 80% power threshold.
